# Duplication and sub-functionalisation characterise diversification of opsin genes in the Lepidoptera

**DOI:** 10.1101/2022.10.31.514481

**Authors:** Muktai Kuwalekar, Riddhi Deshmukh, Saurav Baral, Ajay Padvi, Krushnamegh Kunte

## Abstract

Gene duplication is a vital process for evolutionary innovation. Functional diversification of duplicated genes is best explored in multicopy gene families such as histones, hemoglobin, and opsins. Rhodopsins are photo-sensitive proteins that respond to different wavelengths of light and contribute to diverse visual adaptations across insects. While there are several instances of gene duplications in opsin lineages, the functional diversification of duplicated copies and their ecological significance is properly characterised only in a few insect groups. We examined molecular and structural evolution that underlies diversification and sub-functionalisation of four opsin genes and their duplicated copies across 132 species of the diverse insect order—Lepidoptera. Opsins have largely evolved under purifying selection with few residues showing signs of episodic and pervasive diversifying selection. Although these do not affect overall protein structures of opsins, substitutions in key amino acids in the chromophore-binding pocket of duplicated copies might cause spectral sensitivity shifts leading to sub-functionalisation or neofunctionalisation. Duplicated copies of opsins also exhibit developmental stage-specific expression in *Papilio polytes*, suggesting functional partitioning during development. Together, altered spectral sensitivities owing to key substitutions and differential expression of duplicated copies across developmental stages might enable enhanced colour perception and improved discrimination across wavelengths in this highly visual insect group.

---

Phenotypic and functional diversity across taxa often evolves from a few shared evolutionary and genetic mechanisms, such as co-option and gene duplication. There are numerous examples of adaptations arising as a result of gene duplication (Chen et al. 2008; Kondrashov 2012; Dalla and Dobler 2016; Vandewege et al. 2016 Feb 9). While gene duplication itself is quite common, it is often followed by pseudogenization or nonfunctionalisation resulting in the loss of duplicated copies(Lynch and Conery 2000). Duplicated copies of a gene could undergo neo-functionalisation, where one or more copies may gain an entirely new function (Ohno’s neofunctionalisation), or sub-functionalisation where both copies preserve an element of the ancestral function (DDC-duplication-degeneration-complementation) (Ohno 1970; Force et al. 1999). In both cases, the occurrence of key substitutions in the gene copy is crucial for diversification of functional space. Exploring evolutionary divergence in sequence and structure of duplicated copies, as well as their spatio-temporal dynamics can help us illustrate the role of gene duplication in the expansion of functional space and evolutionary innovation.

Rhodopsins—the primary colour sensing proteins—exhibit an abundant number of paralogs that offer an ideal framework to study functional diversification of duplicated gene copies. Together with the chromophore, opsins form the mechanistic and genetic basis of colour vision across insect orders and thus, also shape intra- and inter-specific interactions and communication. Localized in the microvilli of the photoreceptor cell, they are composed of seven transmembrane domains which form a chromophore binding pocket where retinal-derived chromophore *(11-cis-3-hydroxyretinal* in *Drosophila melanogaster)* is bound to the opsin by a schiffbase linkage (Bownds 1967; Smith and Goldsmith 1990). Incidence of light causes conversion of this chromophore to an *all-trans* state leading to transduction of the light signal and a cascade of events also known as phototransduction (Hardie 2001). Typically, the peak wavelength sensitivity (λ_max_) of opsins depends on the interaction between the chromophore and the opsin protein (Chan et al. 1992; Saito et al. 2019). Mutations in the chromophore binding pockets of opsins can alter their spectral sensitivities and λ_max_, which is often observed in instances of opsin duplications followed by sub-functionalisation or neo-functionalisation (McCulloch et al. 2016; Liénard et al. 2021). Photoreceptor genes have undergone remarkable evolution in insects resulting in an expansion of photoreceptor types and wavelength sensitivities. For example, in a few Diptera, up to six classes of photoreceptors have been reported (Hardie 1986), and several species of Odonata and Hymenoptera can sense a broader range of wavelengths (~350-650 nm) than most other insects (Bybee et al. 2012).

Lepidoptera is an insect order that includes moths and butterflies, which are particularly visual insects and rely heavily on their colour discrimination ability for intra- and inter-specific interactions such as mate choice and predator avoidance. Butterflies diverged from moths nearly 98.3 million years ago (Kawahara et al. 2019), and these groups generally vary in their activity period with moths being largely (but not always) nocturnal and butterflies largely diurnal. Despite the wide spectrum of light environments occupied by moths and butterflies, there are few comparative studies that assess the impact of light environment on the evolution of opsins across this order (Sondhi et al. 2021). In response to varied light environments, Lepidoptera have evolved to possess diverse sets of opsins, which include three commonly known paralogues, i.e., UV opsin λ_max_=300-400 nm), blue or short-wavelength (SW) opsin (λ_max_=400-500 nm), and red or long-wavelength (LW) opsin λ_max_=500-600 nm) (Briscoe 2008). An additional opsin, Rh7 was identified in several insects, which shows poor homology across species and its function is largely unknown. Butterflies and moths have also gained photosensitivity outside the ranges of these three opsins via duplication and neofunctionalisation (Futahashi et al. 2015; Arikawa et al. 2017). For example, genus-specific duplication in the UV opsin in *Heliconius* has resulted in expanded wavelength perception in the UV range due to spectral tuning and is associated with mate choice (Finkbeiner and Briscoe; Briscoe et al. 2010; McCulloch et al. 2016). Previous studies on vision in Lepidoptera have focused on characterization of spectral sensitivities of photoreceptors, spectral tuning mechanisms in different taxa (McCulloch et al.; Wakakuwa et al. 2010) and the effect of diel-niche on vision (Xu et al. 2013; Feuda et al. 2016; Sondhi et al. 2021), the molecular evolution of duplicated opsins, and their role in visual perception remains poorly explored. Moreover, most of these studies address light perception in adults, however, photoreception in larval stages is known to be associated with key adaptations. For instance, larvae of *Biston betularia* are capable of spectrally sensitive extra-ocular photoreception that enables background matching and camouflage (Eacock et al. 2019). More comprehensive studies of visual perception in early stages can help elucidate the mechanistic bases of such adaptations, as well as the temporal dynamics of opsin activity through the life cycle of insects.

We examined the molecular evolution of three well-characterized opsin genes—UV, blue and LW—in Lepidoptera, along with the enigmatic Rh7 opsin. We focused on 132 species of butterflies and moths across 18 families of Lepidoptera. Using gene sequences and developmental transcriptome data we explored the following questions: i) What pattern of molecular and structural evolution do duplicated copies of opsins show? Do they exhibit phylogenetic structure? ii) Do duplicated copies show variations in key residues with a possible involvement in subfunctionalisation or neofunctionalisation by spectral tuning? iii) Do opsins exhibit developmental stage-specific or sexually dimorphic expression across tissues through development? We demonstrate that despite showing conservation at the sequence level, opsin genes exhibit taxon-specific variations in key residues interacting with the chromophore, which could cause wavelength shifts and altered spectral sensitivity. We further show stage-specific partitioning of expression in duplicated opsins in developing *Papilio polytes.*

## RESULTS

### Opsin genes have undergone several lineage-specific duplication events

We obtained opsin sequences from 132 species of Lepidoptera (Dataset S1) and constructed well-supported gene trees for blue, long wavelength (LW), ultra-violet (UV) and Rh7 opsins (Fig. 1). All four gene trees showed a strong pattern of divergence and species relationships that varied across genes (Fig. 1). As previously reported (Briscoe 2001; Briscoe 2008; Feuda et al. 2016), we found several duplication events across Lepidoptera in all opsin genes except for Rh7 (Table S4). However, these duplication events were more common than previously believed. Moreover, in most cases, duplication events were observed across entire tribes or families, and duplicated sequences clustered by copies. We obtained UV opsin sequences from 115 species of moths and butterflies, of which 26 species carried two copies of the gene with a single duplication event (Fig. 1). In addition to the 25 species in the *Heliconius* lineage, *Plutella xylostella*, a nocturnal species of moth, carried two copies of UV opsin. This is surprising given its heavy dependence on olfactory and gustatory stimuli in its habitat (Justus and Mitchell 1996) We obtained sequences for blue opsin in 115 species, of which, 15 butterflies showed duplicated copies while moths carried a single copy (Fig. 1). There were three duplication events, one in Lycaenidae and two in Pieridae. Four species— *Colias erate, Eurema hecabe, Gonepteryx rhamni* and *Phoebis sennae—*carried three copies each of blue opsin. We obtained LW opsin sequences from 122 species and 28 of these showed multiple copies (Fig. 1). The most remarkable bout of gene duplication in LW opsin was observed in Papilionidae, in which *Papilio polytes, P. xuthus* and *P. glaucus* carried six gene copies implying five duplication events. Lineages of both diurnal and nocturnal moths and butterflies exhibited multiple duplication events for LW opsin. For instance, *Callimorpha dominula* (scarlet tiger moth), a diurnal moth showed four copies of LW opsin, while nocturnal species such as *Helicoverpa sp.* belonging to Noctuidae carried two copies each of LW opsin.

**Figure 1.**
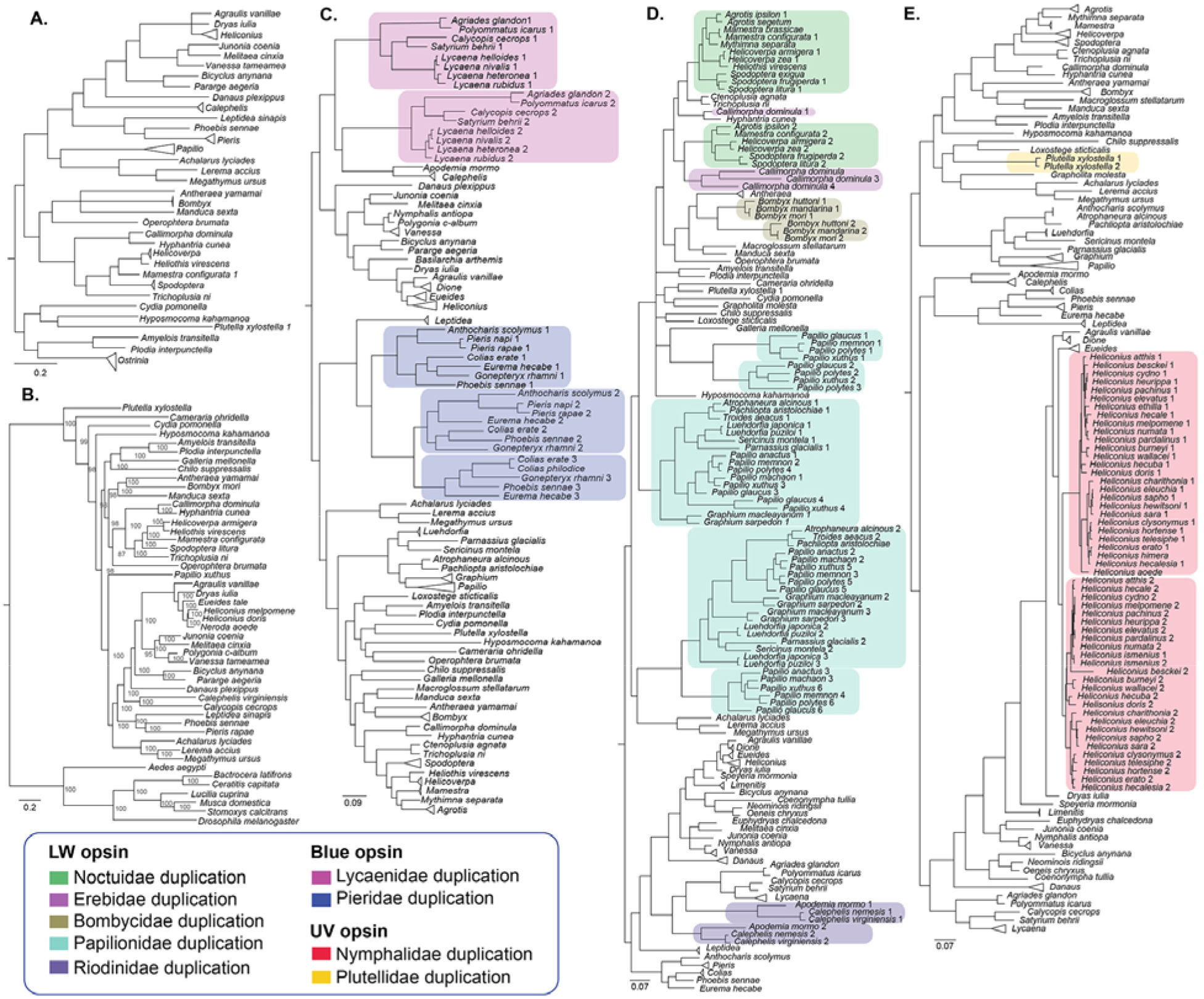
Phylogenetic reconstruction of species tree and gene trees for the four opsin genes. Gene trees were constructed using sequences of the respective opsin genes. The duplication events are colour coded for each of the family. **A.** Genus-level species tree built using two mitochondrial and 11 nuclear genes for the taxa used in this study. **B.** Gene tree for Rh7 gene. **C.** Gene tree for blue opsin gene showing duplication in Lycaenidae and Pieridae. **D.** Gene tree for LW opsin gene showing duplication events in Noctuidae, Erebidae, Bombycidae, Papilionidae and Riodinidae families. **E.** Gene tree for UV opsin gene showing the duplication event in the genus *Heliconius.*

### Opsin genes have evolved under purifying selection with diurnal environments driving diversification across opsins

We plotted site-wise dN, dS and dN/dS for each opsin gene (Fig. 2). Molecular evolution of all four opsin genes was characterized largely by synonymous substitutions, with all sites (with *p*<0.05) showing dN/dS below 1, suggesting a purifying pattern of selection. The mean dN/dS values of opsin genes (UV=0.06, blue=0.06, LW=0.07, Rh7=0.08) were only marginally higher than mean dN/dS of genes used in species-level phylogenetic reconstruction which ranged between 0.006 and 0.04 (Kuwalekar et al. 2020) implying that although opsin genes exhibit a purifying pattern of selection, some amount of diversification has occurred. While gene-wide dS did not differ significantly across opsins, gene-wide dN showed significantly different distribution for blue, UV and Rh7 opsins (Dunn’s test, *p*<0.05) (Fig. 2, Table S6A). None of the sites showed signs of pervasive diversifying selection across all opsin genes and >70% sites showing signatures of pervasive purifying selection (as detected by FEL) (Table S1). Despite the dearth of signatures of diversification, both MEME and BUSTED were able to detect site- and gene-level episodic selection with a considerable number of sites lying in one of the seven transmembrane domains of UV, blue and LW opsins (Table S1, TableS5). In the case of Rh7 opsin, none of the sites experiencing episodic selection were in the domain regions (Table S5). To assess the impact of the activity periods on the evolutionary trajectories of opsins, we partitioned the sequence data into moths and butterflies which are largely nocturnal and diurnal respectively. We found considerable differences in patterns of selection when nocturnal and diurnal species were treated as distinct groups. While dN was significantly different between nocturnal moths and diurnal butterflies for all opsins except Rh7 (Table S6B), dS differed significantly only in UV and Rh7 opsins (Mann-Whitney U test, *p*<0.05) (Table S6B). The mean dN/dS of for diurnal butterflies (ω_UV_=0.0564, ω_Blue_=0.0663, ω_LW_=0.0709) was consistently higher than that for nocturnal moths (ω>_UV_=0.0256, ω_Blue_=0.0318, ω_LW_=0.0352). To further explore this pattern, we performed Contrast-FEL and found a greater number of sites with significantly higher dN/dS (*p*<0.05) diurnal butterflies than in nocturnal moths in all three opsins (Table S2A). Additionally, we used BUSTED and MEME to detect gene-wide and site-wise episodic selection. Consistent with the rate of molecular evolution, we detected more signatures of diversification among diurnal butterflies than nocturnal moths for both gene- and site-level comparisons (Table S2A). It is important to note that poor alignments and low sequence availability for Rh7 may have influenced the selection analysis making it difficult to draw definite conclusions for the evolution of Rh7 in Lepidoptera. We also partitioned the sequence data based climatic zones that the species occupy, i.e., temperate and tropical zones. Across all four opsins, we found very few sites with high dN values (Fig. S1B). The lack of differences in the number of sites undergoing diversifying or purifying selection and absence of variation in the overall dN/dS distribution between these two groups suggests that differences in light conditions due to climatic zone may not be a significant factor in the evolution of opsins (Table S6B and Table S6B).

**Figure 2.**
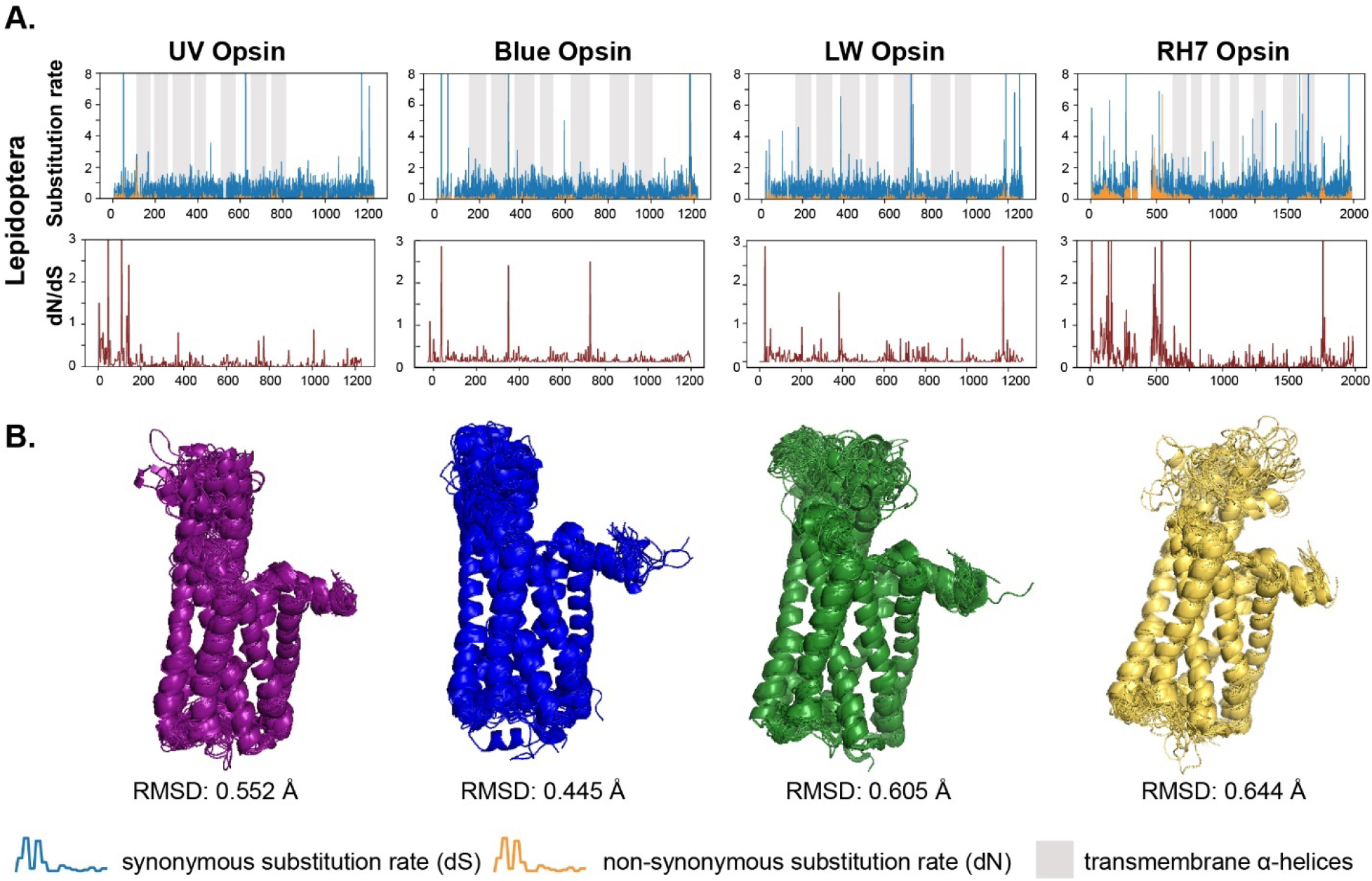
Molecular and structural evolution of opsins in Lepidoptera. **A.** Molecular evolution of opsin genes in Lepidoptera is shown with synonymous (blue) and nonsynonymous substitution rates (orange) for each codon (top panels), and dN/dS ratios for individual codons (lower panels). The seven transmembrane domains for each gene are highlighted in grey. **B.** Structural deviation in the modelled opsin proteins is shown for each opsin with RMSD as a measure of the deviation. Only transmembrane domains were superimposed to calculate RMSD values shown below.

### Opsin proteins show structural conservation across 100 million years of evolution

To measure the structural deviation in protein backbones of opsins, we modelled opsin protein structures and calculated root mean square deviation (RMSD) for each opsin gene. We assessed RMSD values for all Lepidoptera, and separately for groups partitioned across activity periods and climatic zones, by superimposing the domain regions as well as the entire protein model. Across all four opsins, the RMSD values ranged from 0.445 to 0.644 (for domain superimposition), suggesting an overall conserved pattern of structural evolution. The differences in the RMSD values were marginal when structural alignment of entire proteins was compared to that of domain regions only. Rh7 opsin was an exception to this where the RMSD for the entire protein alignment was much higher (2.35) than that for the domain regions (0.6) (Fig. 2B, Fig. S2, Table S3). This indicates that compared to the loop regions of other opsins, Rh7 may have undergone more rapid molecular and structural evolution. Since we consistently observed a diversifying pattern of molecular evolution in diurnal butterflies, we expected structural variation among opsins of diurnal butterflies and nocturnal moths. Interestingly, we did not find differences between protein backbones in any of the opsins among these groups (Table S3), which suggests that the non-synonymous changes at the sequence level are not reflected at the structural level, which is highly conserved. As observed with the patterns of molecular evolution, comparisons between species belonging to temperate and tropical climatic zones revealed no difference in RMSD values (Table S3). Overall, in all opsins but Rh7, lower RMSD values indicate constrained structural evolution.

### Substitutions in the key residues of the chromatophore-binding pocket of opsins may lead to diversification of duplicated copies

We compared amino acid residues that interact with the chromophore across duplicated copies to identify candidates arising from gene duplication with the potential for subfunctionalisation or neofunctionalisation. For example, substitutions resulting in change of polarity at residues 116 S/A and 177 F/Y are crucial for spectral sensitivity of the blue opsin in *Pieris rapae.* (Wakakuwa et al. 2010). We compared amino acid substitutions within a range of 5 Å from the chromophore in the duplicated copies of blue, LW and UV opsin. Blue opsin, which underwent three duplication events in total across Pieridae and Lycaenidae, had seven out of 21 residues within the 5 Å region with substitutions in the duplicated copies (Fig. 3A, Dataset S2). Apart from substitutions at sites 116 and 177, three residues (112A/S, 204I/C, 205F/C/S) showed a change in polarity and two additional residues showed substitutions across duplicates (120I/F, 278A/G) without a change in polarity (Fig. 3A, Dataset S2). Of these seven substitutions, 112, 205, 278 were specific to Pieridae and 204, 120 specific to Lycaenidae (Fig. 3B). Similarly, LW opsin which showed up to six duplicated copies in Papilionidae, had substitutions in six out of 20 sites in the 5 Å region around the chromophore (Fig. 3A, Dataset S2). Interestingly, substitution at 112G/A was also present in diurnal and nocturnal moths (Bombycidae, Erebidae and Noctuidae) but the polarity in this case remained unaltered (Fig 3B). UV opsin had 17 residues within the 5 Å distance. Of these, two showed substitutions without a change in polarity (187T/S/L, 209Y/C), while three sites had substitutions that altered the polarities of these residues (112A/S, 116S/A, 185S/A) (Fig. 3A, Dataset S2). All of these substitutions in UV opsins were specific to Heliconius butterflies (Nymphalidae). Interestingly, one residue, 112, repeatedly showed substitutions in all three opsins with altered polarity in UV (Nymphalidae) and blue opsin (Pieridae) and no alteration in polarity in LW opsin (Papilionidae, Bombycidae, Erebidae and Noctuidae) (Fig. 3B-C). While previous studies have shown an association between change in polarity of residues in the chromophore binding pocket and spectral tuning (Wakakuwa et al. 2010), its mechanism and possible role in Lepidoptera vision is poorly characterized.

**Figure 3.**
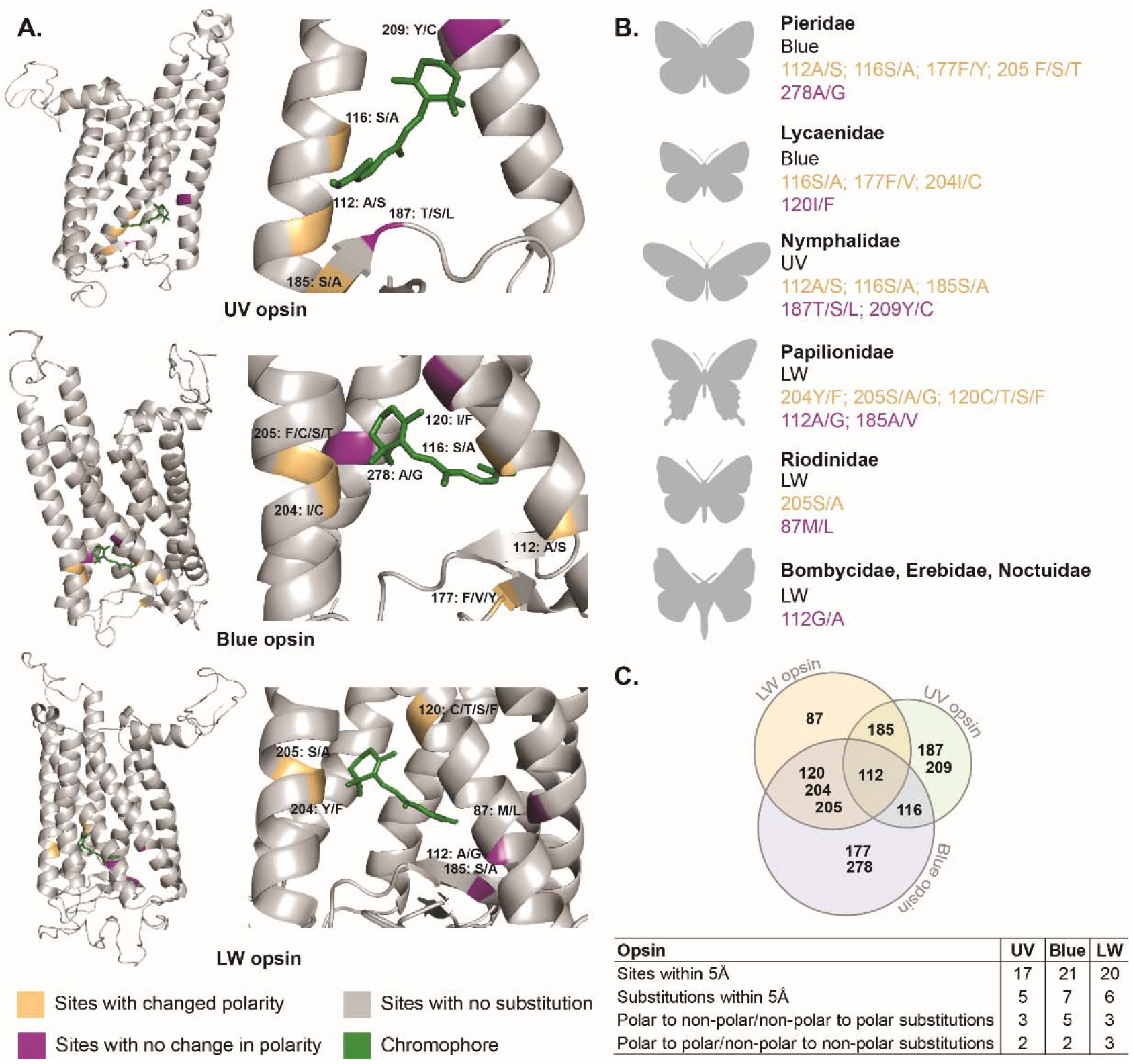
Substitutions in the residues of chromophore-binding pocket potentially involved in spectral tuning. **A.** Modelled protein structures of UV, blue and LW opsins. The chromophore-binding pocket with residues that change the polarity (yellow) or retain the polarity (purple) are shown for each opsin. **B.** Binding pocket residues with substitution in duplicated copy of the opsin gene are shown in a family-wise manner. **C.** Among the binding pocket amino acids, one residue commonly shows substitutions across three opsins while other residues show substitutions in either two or one opsin only.

### Duplicated copies of opsins show distinct expression profiles across developmental stages suggesting functional partitioning

We identified gene sequences of UV, blue, LW and Rh7 opsins in the genome of *P. polytes* using annotations available on NCBI and comparing sequence similarity with well-characterized opsin sequences from other species of Lepidoptera. We identified five copies of LW opsin gene (opsin 1) and single copy each of UV (opsin 2), blue opsin (opsin 3) and Rh7 each, suggesting that LW opsin had undergone one additional duplication event than previously known in *Papilio* (Briscoe 1998; Briscoe 2000). We traced the expression of these eight opsin genes across different developmental stages, tissues, and sexes in *P. polytes* (Fig. 4). Broadly, we identified two groups of LW opsin copies based on their expression patterns*. opsin 1, opsin 1-like 1* and *opsin 1-like 2* formed one group with expression exclusively in late pupal and adult stages. The expression pattern of UV and blue opsin genes was very similar to these copies of LW. The second group comprising *opsin 1-like 3* and *opsin 1-like 4* showed peak expression early during larval development. Rh7 showed a sexually dimorphic expression with peak expression in the body and wing discs of female pre-pupae and little or no expression in pupal and adult stages (Fig. 4A). We also looked for differences in opsin expression between mated and unmated adults, due to the possibility of altered spectral sensitivities during mate choice and oviposition. However, we did not find any difference in opsin expression with change in mating status.

**Figure 4.**
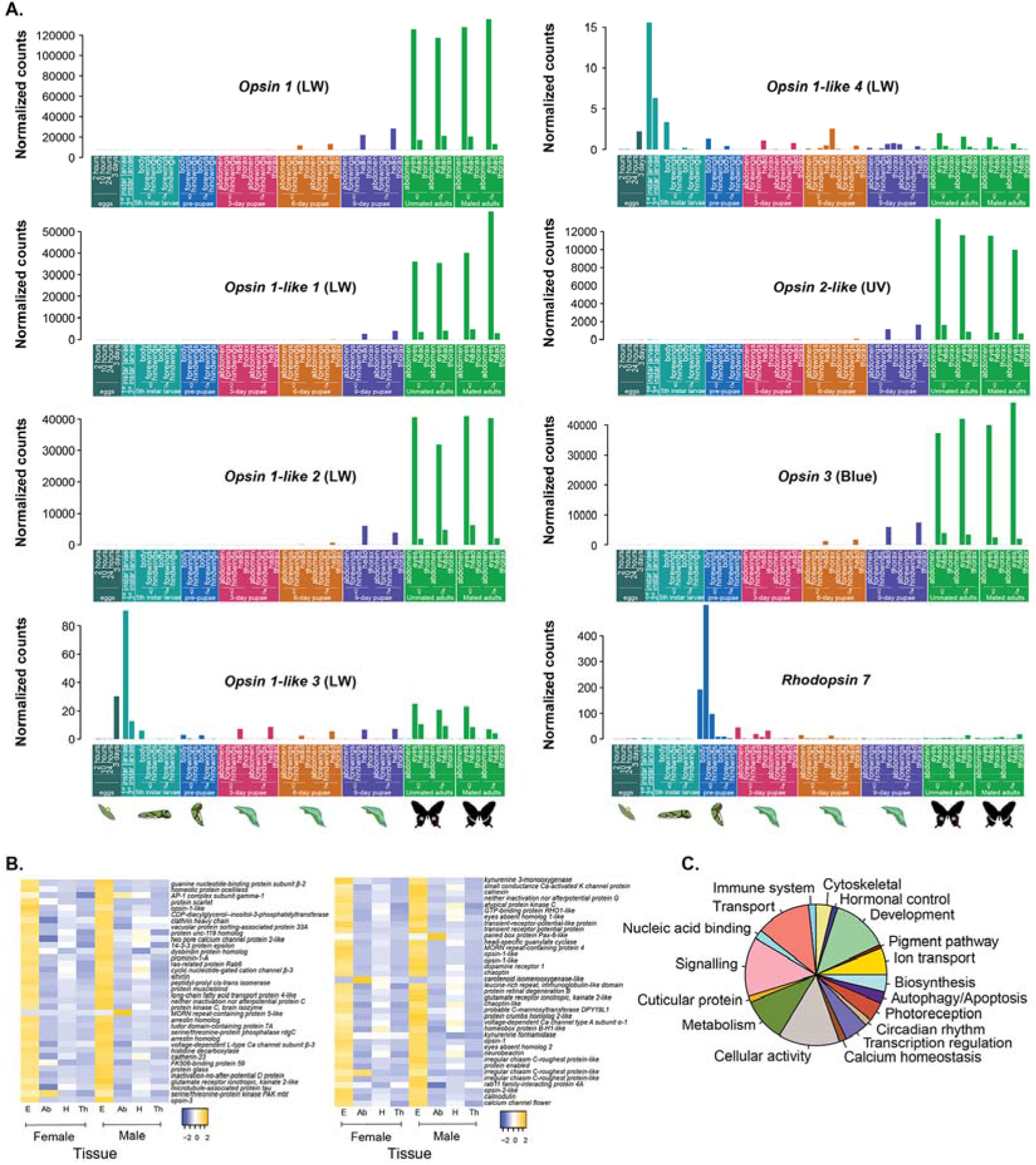
Developmental expression of UV, blue, LW and Rh7 opsin genes in *Papilio polytes.* A. Normalized counts obtained from developmental transcriptome data are plotted for each stage, tissue and sex (beyond 5^th^ instar larvae). The developmental stages are colour-coded and pictorially represented at the bottom. Copies of LW opsin (opsin 1) are sequentially numbered as opsin 1-like genes. **B.** Heatmap for expression of 73 genes involved in vision related pathways. Yellow tiles represent higher expression while blue tiles represent lower expression. **C.** Pie-chart for 803 genes highly expressed in eyes, categorized according to their functions.

We identified 803 genes which showed eye-specific and eye-biased expression in adult *Papilio polytes.* Upon manually annotating these genes for their function, we sorted them into 18 broad categories (Fig. 4C). We found 73 of these 803 genes were involved in photoreception and eye morphogenesis (Table S7). We traced the expression of these 73 genes across tissues (eyes, head, thorax and abdomen) and sexes in adult *Papilio polytes.* Some of these genes, which are primarily involved in eye development, also showed non-ocular expression (Fig. 4B).

## DISCUSSION

Perception of visual signals is a complex process that has evolved over millions of years. The process of phototransduction comprises an intertwined network of pathways which aids in converting light stimuli into electrical signals. Opsins are the primary photoreceptor in this pathway, and also the best studied. In insects, a key component of the diversification and expansion of visual acuity and perception has been the duplication of opsin genes in different lineages. Gene duplication is a well-studied mechanism that facilitates evolution of novel gene functions (Briscoe 2001; Storz et al. 2013; McCulloch et al. 2022). Typically, the most conserved photopigments across insects are those that sense UV, blue and LW (Chang et al. 1996; Townson et al. 1998; Wakakuwa et al. 2005; Arikawa et al. 2017). We identified lineage-specific duplications in three opsin genes (UV, Blue and LW) where the duplicated copies formed clusters based on the sequence homology (Fig. 1). The most extraordinary duplication events were present in LW opsin with three species of *Papilio—P. polytes, P. xuthus, P. glaucus—* which carried six copies of the same. In *P. xuthus* three copies have been well characterized, λm_ax_=515, 530, 575 nm (Arikawa 2003; Briscoe 2008) Similarly, in *P. glaucus* a total of four LW opsin copies have been identified and characterized (Briscoe 1998; Briscoe 2000). We were able to detect the presence of two additional copies with sequence homology to LW opsins. Their presence warrants further investigation to characterize their spectral sensitivity and functional role in insect vision. Blue opsin, on the other hand, seems to have undergone duplication in Pieridae and Lycaenidae. Two copies of short wavelength opsins in *Pieris rapae* have been previously characterized as violet (425 nm) and blue (453 nm) with violet showing sexually dimorphic spectral sensitivity presumably due to spectral filtering in males. Whether such dimorphism is also observed in duplicated copies of blue opsin in other Pieridae and Lycaenidae remains to be seen.

We explored the implications of these rampant duplications by examining key sites that might be involved in subfunctionalisation or neofunctionalisation of duplicated copies. The range of wavelengths that isomerize the chromophore depends on the interaction between amino acids in the chromophore binding pocket of the opsin protein and the chromophore (Chan et al. 1992; Saito et al. 2019). Substitutions in these amino acids that cause a change in polarity of the residue also change the electron distribution in the pi-electron system of the chromophore (Honig et al. 1976; Frentiu et al. 2007). This results in altered sensitivity of the given opsin to various wavelengths and can be crucial for vision in Lepidoptera. We used structural alignments to identify residues within the binding pocket and characterized variation in these residues between duplicated copies. We identified one residue at position 112, that repeatedly shows substitutions across all three opsins. We also identified residues which show opsin-specific substitutions, as well residues that show repeated substitution in two out of three opsins (Fig. 3C). Moreover, we examined these residues in a family-specific manner and identified residues which are family- or lineage-specific, and those that vary between families in Lepidoptera. Further characterization of these key residues and the effect of change in polarity at such sites would help elucidate their role in wavelength perception in different opsins. Considering that a single amino acid substitution can result in adaptative modification (Kojima et al. 2017), studying the cumulative contribution of polar to non-polar and non-polar to polar substitutions is imminent to understand how duplications have given rise to newer colour channels by spectral tuning in Lepidoptera. We also tested these substituted sites for episodic diversifying selection. Although we did not find signatures of diversification directly in these sites, we found several sites adjacent to these key residues that showed episodic diversifying selection (Table S5, Dataset S2). In addition to spectral sensitivity shifts in opsins, wavelength absorption spectrum is also dependent on anatomical and physiological factors such as corneal pigmentation, multilayering, optical waveguides, filtering and spectral organization of ommatidia (van der Kooi et al. 2021). Investigating the interactions of these key components would be instrumental to understand the evolution of visual systems.

We assessed the evolutionary trajectories of four opsins across the insect order Lepidoptera at the molecular and structural level. We also addressed whether light environments in different climate zones and activity periods have played a role in the evolution of opsins, we found that butterflies consistently showed signatures of diversification when compared with moths (as previously shown). While the higher number of duplicated copies in butterflies may contribute to higher dN values to an extent, the consistent pattern of diversification in butterflies may not be solely attributed to duplication. Interestingly, even though we observed gene-level differences in butterflies and moths, such changes were not reflected at the structural level. Such a high degree of structural conservation may suggest that smaller rather than large structural changes, such as substitutions in the amino acid residues of the binding pocket may play a larger role in the diversification.

We explored the spatio-temporal activity of duplicated opsins by tracing the expression of five copies of LW opsins and a single copy of blue, UV and Rh7 opsin, through developmental stages of *P. polytes* using a developmental transcriptome dataset. We found partitioning in the expression of LW opsins where two copies were expressed in larval stages and three copies were expressed exclusively in adult stages. Whether these LW opsins play a role in ocular and/or extraocular visual perception in larval stages, remains to be explored. Interestingly, the relatively poorly characterized opsin, Rh7 showed a sexually dimorphic expression. It is possible that the expression of Rh7 may be transient and our sampling may have captured this in females, but missed it in males. However, further investigation of the role of Rh7 in developing Lepidoptera may help explain this curious expression pattern. We also traced expression of 73 genes that show eye-specific and vision related function. Some of these showed expression in abdomen and thorax in adults, possibly suggesting a role in extra-ocular vision. Interestingly, extraocular vision plays a role in background matching in peppered moth larvae (Eacock et al. 2019) which also occurs in *Papilio polytes* pupae, which show green or brown colouration based on whether they pupate on leaves or twigs respectively. The decision of background matching is usually made by wandering larvae just before pupation, based on a variety of cues (Yoda et al. 2020). Further experimental validation might help determine which set of opsins (LW or Rh7) play a role in background matching in *Papilio polytes.* Genetic manipulation of individual opsins and its effect on visual perception in *Papilio* butterflies may help elucidate their ecological and behavioral relevance.

We demonstrate that opsin genes in Lepidoptera show a pattern of conservation and yet, they exhibit extensive gene duplication and diversification. We highlight key residues that could potentially enable diversification of duplicated opsins in the Lepidoptera. Our work provides a comparative assessment between genes and species to understand patterns of conservation and variation in the molecular and structural evolution of opsins. Our work also provides insights into partitioning of opsin expression, and possibly function, across developing *P. polytes.* Further exploration of larval and extraocular vision in holometabolous insects can help us understand dynamic vision systems through metamorphosis.

## Acknowledgments

We thank Vinod Shankar for assistance in downloading and analysing genome sequences, NCBS Sequencing Facility for transcriptome sequencing, and NCBS Greenhouse Facility for butterfly breeding facilities.

## Author contributions

SB and KK designed the study. SB and AP downloaded and curated the sequence data. SB performed the gene and species tree construction. MK performed molecular evolution analysis. AP performed the structural evolution analysis. RD performed the developmental transcriptome sequencing. MK and RD analysed the developmental transcriptome data. MK and RD prepared the figures and wrote the manuscript. KK conceived and directed the project. All authors contributed to the article and approved the submitted version.

## Competing interests

The authors declare no competing interests.

## Additional Information

***Supplementary Information*** is available for this paper.

### Funding

This work was partially funded by an NCBS Research Grant to KK, a CSIR Shyama Prasad Mukherjee Fellowship to RD, and NCBS Student Fellowship to SB.

### Data Availability Statement

The raw RNA seq data are deposited in the NCBI SRA database (BioProject PRJNA634605; Accession nos. SAMN15001929 SAMN15002046to; https://dataview.ncbi.nlm.nih.gov/object/PRJNA634605?reviewer=mf4q72fcfvq67spjf5gl1sduop)

***Correspondence and requests for materials*** should be addressed to.

## METHODS

### Data mining, multiple sequence alignment and identification of duplicates

We downloaded whole genome sequences of 90 species of Lepidoptera (moths and butterflies) from GenBank, LepBase, and GigaDB. We also downloaded available opsin sequences for UV (rhodopsin 4), Blue (rhodopsin 5), LW (rhodopsin 6) and rhodopsin 7. We performed exon-wise local NCBI BLAST+ with tBLASTn to locate gene sequences for all four opsins within the downloaded genomes. For UV, blue, LW and RH7, we were able to obtain sequences for 115, 115, 122 and 62 species out of which 26, 15, 28 and 0 species carried more than one copy of the respective opsin. We set a cut off of 80 percent similarity to identify the given sequence as a duplicate. We performed multiple sequence alignments with PRANK version v150803 (Löytynoja and Goldman 2010). To assess effect of light environment in the daily activity period and climate zones on the evolution of opsins, we partitioned sequences into moths and butterflies and temperate and tropical Lepidoptera. We were able to extract sequences for not more than 1 −2 species of nocturnal butterflies or diurnal moths each. Thus, the sequence data was partitioned into butterflies and moths rather than diurnal and nocturnal. We used MUSCLE aligner on MEGA X with codon alignment option to align these partitioned sequences as PRANK introduced additional gaps during alignment (Kumar et al. 2018). We determined the sequence corresponding to seven transmembrane domains within each opsin gene using CD-Search Tool and Conserved Domain Database (CDD) (Marchler-Bauer et al. 2012).

### Phylogenetic analysis and construction of gene trees

We constructed a species level phylogeny with 41 species of Lepidoptera and seven outgroups from Diptera. We used two mitochondrial *(cytochrome c oxidase I, acetyl-CoA acetyltransferase)* and 11 nuclear genes *(elongation factor 1-alpha, wingless, ribosomal protein S5, ribosomal protein S2, isocitrate dehydrogenase, glyceraldehyde-3-phosphate dehydrogenase, malate dehydrogenase, catalase, CAD, ribosomal protein S27a/hairy cell leukemia, dopa-decarboxylase)* as phylogenetic markers (Wahlberg and Wheat 2008). We aligned these 13 gene sequences with codon aligner tool of PRANK version v150803 and computed the best partition scheme and evolution models for both, marker alignments and individual opsin gene alignments using Partition Finder 2.1.1 (Lanfear R et al. 2012). We selected greedy algorithm and MrBayes model option, whereas, to compare the best-fit models, we chose BIC (Bayesian information criterion). We set a split frequency cut-off of 0.01 and used the remaining trees to construct a consensus tree. The same pipeline was followed to construct gene trees for all four opsin genes.

### Computation of synonymous (dS) and non-synonymous (dN) substitution rates

We estimated sitewise synonymous and non-synonymous substitution rates for each opsin gene using the PRANK alignments. We used fixed effect likelihood (FEL) which calculates maximum-likelihood to determine synonymous (dS) and non-synonymous substitution (dS) rates for each site (Kosakovsky Pond and Frost 2005). We identified sites under pervasive diversification (dN/dS > 1) or purifying selection (dN/dS < 1) with FEL. We plotted dN/dS values for all Lepidoptera using custom python scripts and compared the evolutionary rates of moths and butterflies, and temperate and tropical Lepidoptera. We performed Mann-Whitney U tests to compute p-values for pairwise dN and dS comparisons between moths and butterflies, and temperate and tropical Lepidoptera. For comparing dN and dS of each opsin with the other, we performed Kruskal-Wallis test and Dunn’s test. We used Bonferroni correction to adjust p-values for multiple comparisons. To compare diversifying and purifying patterns between butterflies and moths, we performed Contrast-FEL which tests for differences in selective pressures (Kosakovsky Pond et al. 2020). We performed FEL, MEME, BUSTED and Contrast-FEL on the Datamonkey Adaptive Evolution Server (Murrell et al. 2012; Smith et al. 2015; Weaver et al. 2018) and processed the data using custom scripts.

### Structural alignments and homology modelling

We used ExPASy translate tool to translate gene sequences of each opsin for all Lepidoptera species used. We performed structural alignments and visualized these using PyMOL (2.0.6). We modeled entire protein sequences for each species using Phyre2. We refined these models using GalaxyRefine tool for stearic hinderance on the Galaxy WEB server. Using the confidence score from Phyre2 and refinement scores, we chose one structure from one species as a base model for each opsin, for structural superimposition of all the Lepidoptera protein models in PDB format. We calculated RMSD (Root Mean Square Deviation) from these superimposed models for all four opsins separately. We then partitioned our sequences into butterflies and moths, temperate and tropical Lepidoptera and followed the same pipeline to calculate RMSD for these groups separately. To assess spectral tuning in duplicated copies of blue, LW and UV opsin, we focused on the amino acid residues within a 5 Å radius of the chromophore binding site. We used squid rhodopsin from PDB as a reference to align our modeled structures. We recorded the changes in residues within 5 Å radius across duplicated copies of all opsins.

### Sample collection, RNA extraction and transcriptome sequencing for quantification of the expression

We maintained *Papilio polytes* populations from wild-caught mated females in a greenhouse at 28±4°C. We raised larvae on lemon *(Citrus sp.)* and curry plants *(Murraya koenigii)* and maintained adults on Birds Choice™ butterfly nectar. We sampled several developmental stages from this population for RNA extraction and sequencing. We also separately processed different tissues at several stages for the same. We collected eggs post-oviposition at 2, 10 and 24 hours and at 3 days, and pooled five eggs for each sample to obtain sufficient quantity of RNA. We collected 1^st^, 3^rd^ and 5^th^ instar larvae and used gutted bodies of 1^st^ and 3^rd^ instar larvae, while we dissected 5^th^ instar larvae to separate forewings and hindwings from the gutted body. We dissected 3-, 6- and 9-day old pupae to separate forewings, hindwings, abdomen, thorax and head. To test if mating status has an effect on the gene expression of opsins, we collected mated and unmated adults and dissected them to separate abdomen, eyes, head and thorax. We sampled triplicates of eggs, 1^st^ and 3^rd^ instar larvae, duplicates of 5^th^ instar larvae, pre-pupae and pupae for each sex and quadruplets of adults for each mating status and sex.

We used TRIzol™ to preserve collected samples and used chloroform-isopropanol-based extraction method for RNA extraction. To prepare libraries, we used TruSeq^®^ RNA Sample Preparation Kit v2 and performed Qubit fluorometric quantification. We checked library profiles with Bioanalyzer. To sequence the transcriptome, we performed 2×100 PE runs on Illumina HiSeq 2500 and obtained ~20 million reads per sample. We checked the quality of the reads and appropriately trimmed them using FastQC and Trimmomatic respectively. We aligned these processed reads to the *P. polytes* reference genome (Nishikawa et al. 2015) using STAR aligner for eggs, larvae and pupae and HISAT2 aligner for adults. We obtained raw counts using HTSeq and further analyzed and plotted gene expression for all opsin copies using an edgeR pipeline. In addition to opsin genes, we identified genes involved in vision-related pathways from literature. We plotted expression of these genes along with genes that are highly expressed in eyes.

## SUPPLEMENTARY TABLES AND FIGURES

**Table S1.**
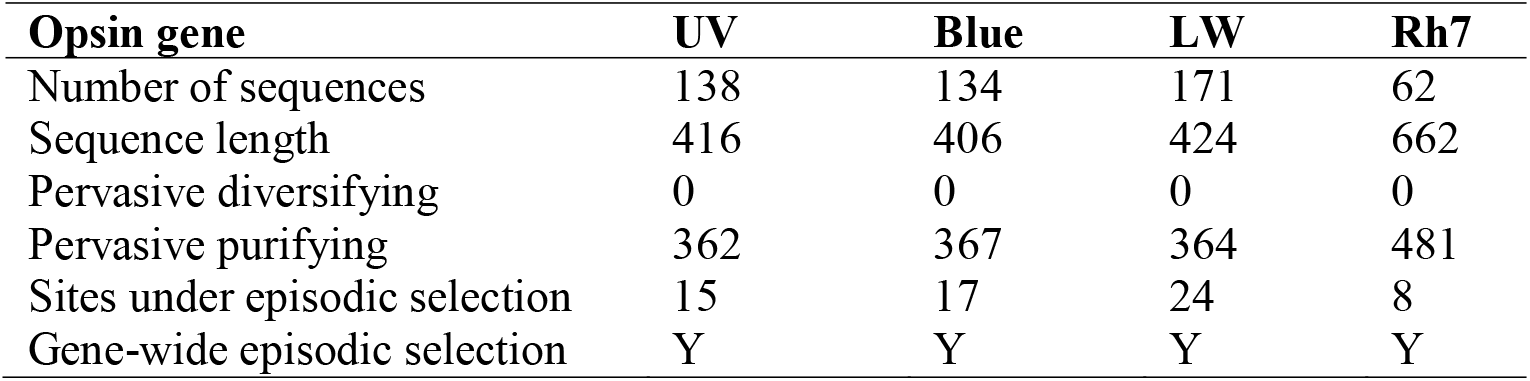
Molecular evolution of opsin genes. A summary of site-wise and gene-level selection analysis of opsin genes in Lepidoptera.

| **Opsin gene** | **UV** | **Blue** | **LW** | **Rh7** |
| --- | --- | --- | --- | --- |
| Number of sequences | 138 | 134 | 171 | 62 |
| Sequence length | 416 | 406 | 424 | 662 |
| Pervasive diversifying | 0 | 0 | 0 | 0 |
| Pervasive purifying | 362 | 367 | 364 | 481 |
| Sites under episodic selection | 15 | 17 | 24 | 8 |
| Gene-wide episodic selection | Y | Y | Y | Y |

**Table S2.** Comparative molecular evolution of opsin genes. A. A summary of comparative selection analysis performed on gene sequences of butterflies and moths separately, and B. temperate and tropical Lepidoptera separately. Significance of *p*<0.05 is indicated with *.

**A.**
| Opsin gene | Blue |  | LW |  | UV |  | RH7 |  |
| --- | --- | --- | --- | --- | --- | --- | --- | --- |
|  | Moths | Butterflies | Moths | Butterflies | Moths | Butterflies | Moths | Butterflies |
| Number of sequences | 30 | 103 | 44 | 127 | 26 | 111 | 22 | 38 |
| Number of sequences used | 30 | 103 | 44 | 126 | 26 | 108 | 22 | 38 |
| Sequence length | 390 | 387 | 387 | 391 | 379 | 390 | 614 | 597 |
| Sites undergoing diversifying selection | 0 | 0 | 0 | 0 | 0 | 0 | 1 | 0 |
| Sites undergoing purifying selection | 321 | 355 | 332 | 351 | 315 | 350 | 414 | 422 |
| Sites undergoing episodic positive selection | 6 | 18 | 7 | 23 | 2 | 19 | 5 | 2 |
| Gene-wide episodic selection | N | Y | Y | Y | N | Y | Y | N |
| Sites with high omega | 9 | 42 | 10 | 58 | 20 | 28 | 42 | 19 |

**B.**
| Opsin gene | Blue |  | LW |  | UV |  | RH7 |  |
| --- | --- | --- | --- | --- | --- | --- | --- | --- |
|  | Temperate | Tropical | Temperate | Tropical | Temperate | Tropical | Temperate | Tropical |
| Number of Sequences | 48 | 66 | 55 | 90 | 38 | 89 | 14 | 39 |
| Number of sequences in used | 48 | 65 | 55 | 89 | 37 | 85 | 14 | 38 |
| Sequence length | 390 | 392 | 393 | 386 | 389 | 391 | 603 | 623 |
| Sites undergoing diversifying selection | 0 | 0 | 0 | 0 | 0 | 0 | 0 | 0 |
| Sites undergoing purifying selection | 349 | 336 | 345 | 350 | 338 | 343 | 384 | 429 |
| Sites undergoing episodic positive selection | 9 | 9 | 16 | 13 | 3 | 15 | 2 | 4 |
| Gene-wide episodic selection | Y | Y | Y | Y | Y | Y | N | Y |

**Table S3.** Group-wise comparison of RMSD values for structural resemblance of opsin proteins. Based on refinement and confidence scores, one structure was used as the base model for superimposition of modelled proteins. RMSD was used as a measure of structural deviation between protein backbones of opsins and is shown for superimposition of entire protein structures as well as the functional domains. RMSD was also calculated separately for butterflies, moths as well as temperate and tropical Lepidoptera for all opsin protein models.

| **Opsin** | **Lepidoptera** | **Butterflies** | **Moths** | **Temperate** | **Tropical** |
| --- | --- | --- | --- | --- | --- |
| Blue - entire protein | 0.6993 | 0.7181 | 0.6151 | 0.692 | 0.653 |
| Blue - domain region | 0.552 | 0.684 | 0.509 | 0.549 | 0.519 |
| LW - entire protein | 0.5008 | 0.5058 | 0.3872 | 0.617 | 0.508 |
| LW - domain region | 0.4456 | 0.4518 | 0.3587 | 0.499 | 0.4546 |
| UV - entire protein | 0.6993 | 0.71854 | 0.5683 | 0.611 | 0.725 |
| UV - domain region | 0.6056 | 0.6117 | 0.5522 | 0.5125 | 0.6587 |
| Rh7 - entire protein | 2.3539 | 3.809 | 2.2577 | 2.24 | 3.425 |
| Rh7 - domain region | 0.6443 | 0.8865 | 0.6530 | 0.6635 | 0.789 |

**Table S4.** Species-wise copies of opsin genes.

| **Species** | **Number of copies** |  |  |  |
| --- | --- | --- | --- | --- |
|  | **Blue** | **UV** | **LW** | **RH7** |
| *Achalarus lyciades* | 1 | 1 | 1 | 1 |
| *Agraulis vanillae* | 1 | 1 | 1 | 1 |
| *Agriades glandon* | 2 | 1 | 1 | 0 |
| *Agrotis Ipsilon* | 1 | 1 | 2 | 0 |
| *Agrotis segetum* | 1 | 1 | 1 | 0 |
| *Amyelois transitella* | 1 | 1 | 1 | 1 |
| *Antheraea pernyi* | 0 | 0 | 1 | 0 |
| *Antheraea yamamai* | 1 | 1 | 1 | 1 |
| *Anthocharis scolymus* | 2 | 1 | 1 | 0 |
| *Apodemia mormo* | 1 | 1 | 2 | 0 |
| *Atrophaneura alcinous* | 1 | 1 | 2 | 0 |
| *Basilarchia arthemis* | 1 | 0 | 0 | 0 |
| *Bicyclus anynana* | 1 | 1 | 1 | 1 |
| *Bombyx huttoni* | 1 | 1 | 2 | 1 |
| *Bombyx mandarina* | 1 | 1 | 2 | 1 |
| *Bombyx mori* | 1 | 0 | 2 | 1 |
| *Calephelis nemesis* | 1 | 1 | 2 | 1 |
| *Calephelis virginiensis* | 1 | 1 | 2 | 1 |
| *Callimorpha dominula* | 1 | 1 | 4 | 1 |
| *Calycopis cecrops* | 2 | 1 | 1 | 0 |
| *Cameraria ohridella* | 1 | 0 | 1 | 0 |
| *Chilo suppressalis* | 1 | 1 | 1 | 0 |
| *Coenonympha tullia* | 0 | 1 | 1 | 0 |
| *Colias erate* | 3 | 1 | 1 | 0 |
| *Colias philodice* | 1 | 1 | 1 | 0 |
| *Ctenoplusia agnata* | 1 | 1 | 1 | 0 |
| *Cydia pomonella* | 1 | 0 | 1 | 1 |
| *Danaus gilippus* | 0 | 1 | 1 | 0 |
| *Danaus plexippus* | 1 | 1 | 1 | 1 |
| *Dione junio* | 1 | 1 | 1 | 0 |
| *Dione moneta* | 1 | 1 | 1 | 0 |
| *Dryas iulia* | 1 | 1 | 1 | 1 |
| *Eueides aliphera* | 1 | 1 | 1 | 0 |
| *Eueides isabella* | 1 | 1 | 1 | 0 |
| *Eueides lineata* | 1 | 1 | 1 | 0 |
| *Eueides procula* | 1 | 1 | 1 | 0 |
| *Eueides vibilia* | 1 | 1 | 1 | 0 |
| *Euphydryas chalcedona* | 0 | 1 | 1 | 0 |
| *Eurema hecabe* | 3 | 1 | 1 | 0 |
| *Galleria mellonella* | 1 | 0 | 1 | 0 |
| *Gonepteryx rhamni* | 3 | 0 | 0 | 0 |
| *Graphium macleayanum* | 1 | 1 | 3 | 0 |
| *Graphium sarpedon* | 1 | 1 | 3 | 0 |
| *Grapholita molesta* | 0 | 1 | 1 | 0 |
| *Heliconius atthis* | 1 | 2 | 1 | 0 |
| *Heliconius besckei* | 1 | 2 | 1 | 1 |
| *Heliconius burneyi* | 1 | 2 | 1 | 1 |
| *Heliconius charithonia* | 1 | 2 | 1 | 0 |
| *Heliconius clysonymus* | 1 | 2 | 1 | 0 |
| *Heliconius cydno* | 1 | 2 | 1 | 1 |
| *Heliconius eleuchia* | 1 | 2 | 1 | 0 |
| *Heliconius elevatus* | 1 | 2 | 1 | 1 |
| *Heliconius erato* | 1 | 2 | 1 | 1 |
| *Heliconius ethilla* | 1 | 1 | 1 | 1 |
| *Heliconius hecale* | 1 | 2 | 1 | 1 |
| *Heliconius hecalesia* | 1 | 2 | 0 | 1 |
| *Heliconius hecuba* | 1 | 2 | 1 | 0 |
| *Heliconius heurippa* | 0 | 2 | 0 | 1 |
| *Heliconius hewitsoni* | 1 | 2 | 1 | 0 |
| *Heliconius hierax* | 0 | 0 | 1 | 0 |
| *Heliconius himera* | 1 | 1 | 1 | 1 |
| *Heliconius hortense* | 1 | 2 | 1 | 0 |
| *Heliconius ismenius* | 1 | 2 | 1 | 1 |
| *Heliconius melpomene* | 1 | 2 | 1 | 1 |
| *Heliconius numata* | 1 | 2 | 1 | 1 |
| *Heliconius pachinus* | 1 | 2 | 1 | 0 |
| *Heliconius pardalinus* | 1 | 2 | 1 | 1 |
| *Heliconius sapho* | 1 | 2 | 1 | 0 |
| *Heliconius sara* | 1 | 2 | 1 | 1 |
| *Heliconius telesiphe* | 1 | 2 | 1 | 1 |
| *Heliconius timareta* | 1 | 0 | 1 | 0 |
| *Heliconius wallacei* | 1 | 2 | 0 | 0 |
| *Heliconius xanthocles* | 0 | 0 | 1 | 0 |
| *Helicoverpa armigera* | 1 | 1 | 2 | 1 |
| *Helicoverpa zea* | 1 | 1 | 2 | 1 |
| *Heliothis virescens* | 1 | 1 | 1 | 1 |
| *Hyphantria cunea* | 1 | 1 | 1 | 1 |
| *Hypsmocoma kahamanoa* | 1 | 1 | 1 | 1 |
| *Junonia coenia* | 1 | 1 | 1 | 1 |
| *Laparus doris* | 1 | 2 | 1 | 1 |
| *Leptidea amurensis* | 1 | 1 | 1 | 0 |
| *Leptidea sinapis* | 1 | 1 | 1 | 1 |
| *Lerema accius* | 1 | 1 | 1 | 1 |
| *Limenitis arthemis* | 0 | 1 | 1 | 0 |
| *Limenitis weidemeyerii* | 0 | 1 | 1 | 0 |
| *Loxostege sticticalis* | 1 | 1 | 1 | 0 |
| *Luehdorfia japonica* | 1 | 1 | 3 | 0 |
| *Luehdorfia puziloi* | 1 | 1 | 3 | 0 |
| *Lycaena helloides* | 2 | 1 | 1 | 0 |
| *Lycaena heteronea* | 2 | 1 | 1 | 0 |
| *Lycaena nivalis* | 2 | 1 | 1 | 0 |
| *Lycaena rubidus* | 2 | 1 | 1 | 0 |
| *Macroglossum stellatarum* | 1 | 1 | 1 | 0 |
| *Mamestra brassicae* | 1 | 1 | 1 | 0 |
| *Mamestra configurata* | 1 | 1 | 2 | 1 |
| *Manduca sexta* | 1 | 1 | 1 | 1 |
| *Megathymus ursus* | 1 | 1 | 1 | 1 |
| *Melitaea cinxia* | 1 | 0 | 1 | 1 |
| *Mythimna separata* | 1 | 1 | 1 | 0 |
| *Neruda aoede* | 1 | 1 | 0 | 1 |
| *Neominois ridingsii* | 0 | 1 | 1 | 0 |
| *Nymphalis antiopa* | 1 | 1 | 1 | 0 |
| *Oeneis chryxus* | 0 | 1 | 1 | 0 |
| *Operophtera brumata* | 1 | 0 | 1 | 1 |
| *Ostrinia furnacalis* | 0 | 0 | 0 | 1 |
| *Ostrinia scapulalis* | 0 | 0 | 0 | 1 |
| *Pachliopta aristolochiae* | 1 | 1 | 2 | 0 |
| *Papilio anactus* | 1 | 1 | 3 | 0 |
| *Papilio glaucus* | 1 | 1 | 6 | 1 |
| *Papilio machaon* | 1 | 1 | 3 | 1 |
| *Papilio memnon* | 0 | 1 | 4 | 1 |
| *Papilio polytes* | 1 | 1 | 6 | 1 |
| *Papilio xuthus* | 1 | 1 | 6 | 1 |
| *Pararge aegeria* | 1 | 0 | 0 | 1 |
| *Parnassius glacialis* | 1 | 1 | 2 | 0 |
| *Phoebis sennae* | 3 | 1 | 1 | 1 |
| *Pieris napi* | 2 | 1 | 1 | 1 |
| *Pieris rapae* | 2 | 1 | 1 | 1 |
| *Plodia interpunctella* | 1 | 1 | 1 | 1 |
| *Plutella xylostella* | 1 | 2 | 1 | 1 |
| *Polygonia c-album* | 1 | 0 | 0 | 0 |
| *Polyommatus icarus* | 2 | 1 | 1 | 0 |
| *Satyrrium behrii* | 2 | 1 | 1 | 0 |
| *Sericinus montela* | 1 | 1 | 2 | 0 |
| *Speyeria Mormonia* | 0 | 1 | 1 | 0 |
| *Spodoptera exigua* | 1 | 1 | 1 | 0 |
| *Spodoptera frugiperda* | 1 | 1 | 2 | 1 |
| *Spodoptera litura* | 1 | 1 | 2 | 1 |
| *Trichoplusia ni* | 1 | 1 | 1 | 1 |
| *Troides Aeacus* | 0 | 0 | 2 | 0 |
| *Vanessa cardui* | 1 | 1 | 1 | 0 |
| *Vanessa tameamea* | 1 | 1 | 1 | 1 |
| **Number of sequences** | **134** | **141** | **172** | **62** |

**Table S5.** Sites undergoing episodic selection in opsin genes.

| **Opsin** | **Sites (Amino acid)** | **Codon** | **Site in polytes**<br>**(Amino acid)** | **Codon_polytes** | **Domain** |
| --- | --- | --- | --- | --- | --- |
| UV | 11 | ACW | 4 | CCT | Non-domain |
| UV | 16 | AAT | 9 | AAT | Non-domain |
| UV | 35 | TCT | 24 | AGA | Non-domain |
| UV | 36 | AGC | 25 | TCG | Non-domain |
| UV | 90 | TGC | 66 | GTG | TM1 |
| UV | 123 | AAT | 99 | CAG | TM2 |
| UV | 135 | GCA | 111 | GCG | Non-domain |
| UV | 169 | GCG | 145 | GGA | TM3 |
| UV | 186 | TCT | 157 | TCT | TM3 |
| UV | 207 | TGT | 178 | TGC | TM4 |
| UV | 258 | AGC | 229 | GTC | TM5 |
| UV | 260 | GTA | 231 | ATT | TM5 |
| UV | 264 | TGT | 235 | ACC | TM5 |
| UV | 316 | CTC | 287 | CTC | TM6 |
| UV | 322 | TTT | 293 | TAC | TM6 |
| Blue | 40 | TCC | 20 | TCA | Non-domain |
| Blue | 74 | TGG | 54 | TAC | Non-domain |
| Blue | 77 | GTC | 57 | ATA | TM1 |
| Blue | 133 | ATC | 113 | GTC | TM2 |
| Blue | 142 | GGC | 122 | GGG | Non-domain |
| Blue | 185 | AAC | 165 | AAC | Non-domain |
| Blue | 208 | GCC | 188 | GCC | Non-domain |
| Blue | 237 | GTG | 217 | GTG | TM5 |
| Blue | 239 | GTC | 219 | GTC | TM5 |
| Blue | 241 | TGC | 221 | TGT | TM5 |
| Blue | 248 | TGC | 228 | GCT | TM5 |
| Blue | 273 | AAA | 253 | CAA | Non-domain |
| Blue | 287 | AAC | 267 | AAC | Non-domain |
| Blue | 320 | GTC | 300 | GTC | TM6 |
| Blue | 323 | ACC | 303 | ACC | TM6 |
| Blue | 344 | GCC | 324 | TGC | TM7 |
| Blue | 378 | CCG | 358 | CAG | Non-domain |
| LW |  | ACA | 4 | GAC | Non-domain |
| LW | 22 | CCC | 8 | CCA | Non-domain |
| LW | 88 | GTT | 65 | GTT | TM1 |
| LW | 104 | TTC | 81 | TTC | TM1 |
| LW | 105 | ACA | 82 | ACA | TM1 |
| LW | 108 | AAG | 85 | AAG | Non-domain |
| LW | 130 | TKC | 107 | ACA | TM2 |
| LW | 131 | ATG | 108 | TGC | TM2 |
| LW | 147 | TTC | 124 | TTT | Non-domain |
| LW | 202 | GGA | 178 | GCT | TM4 |
| LW | 205 | GTT | 181 | GTC | TM4 |
| LW | 213 | GCA | 189 | GCA | TM4 |
| LW | 238 | GAC | 214 | GAT | Non-domain |
| LW | 249 | CGC | 218 | CGA | TM5 |
| LW | 250 | AGC | 219 | AGC | TM5 |
| LW | 258 | TTC | 227 | TTC | TM5 |
| LW | 259 | TGC | 228 | TGC | TM5 |
| LW | 272 | TTC | 241 | TTC | TM5 |
| LW | 302 | GCG | 271 | GCG | Non-domain |
| LW | 343 | GCG | 310 | GCT | Non-domain |
| LW | 348 | CTT | 315 | CTG | TM7 |
| LW | 362 | GTC | 329 | GTT | TM7 |
| Rh7 | 7 | CAA | 3 | CTT | Non-domain |
| Rh7 | 110 | AGA | 98 | TTG | Non-domain |
| Rh7 | 111 | CCA | 99 | CAT | Non-domain |
| Rh7 | 123 | GTG | 108 | TCA | Non-domain |
| Rh7 | 539 | ATT | 508 | ACT | Non-domain |
| Rh7 | 547 | --- | --- | --- | Non-domain |
| Rh7 | 567 | TC- | --- | --- | Non-domain |
| Rh7 | 606 | ACG | 570 | GCG | Non-domain |

**Table S6.**
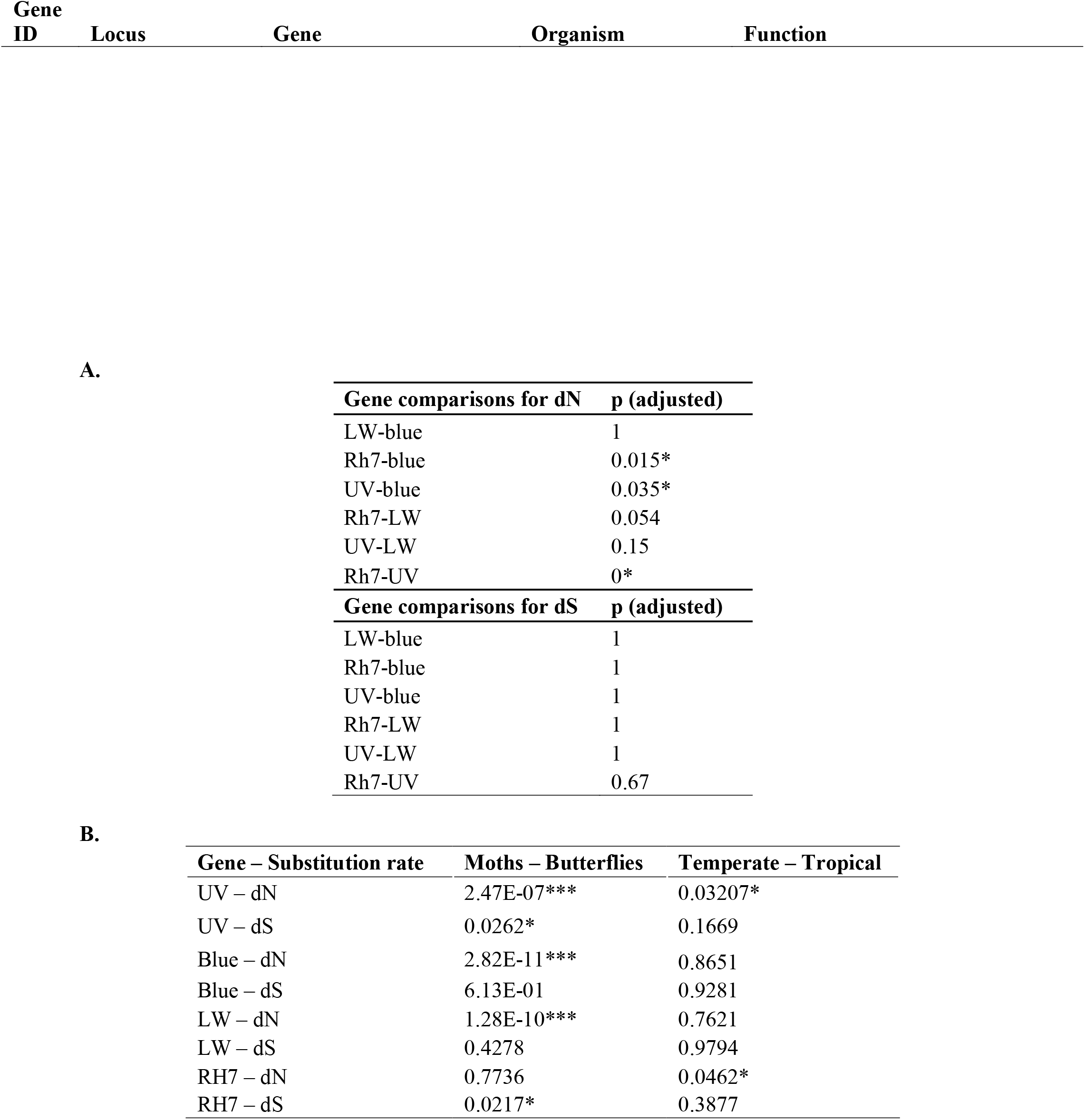
*p*-values for rates of molecular evolution. **A.** *p*-values for pair-wise comparison of dN and dS values between different opsins across Lepidoptera, and **B.** between butterflies and moths, and Lepidoptera from different climate zones.

**Table S7.**
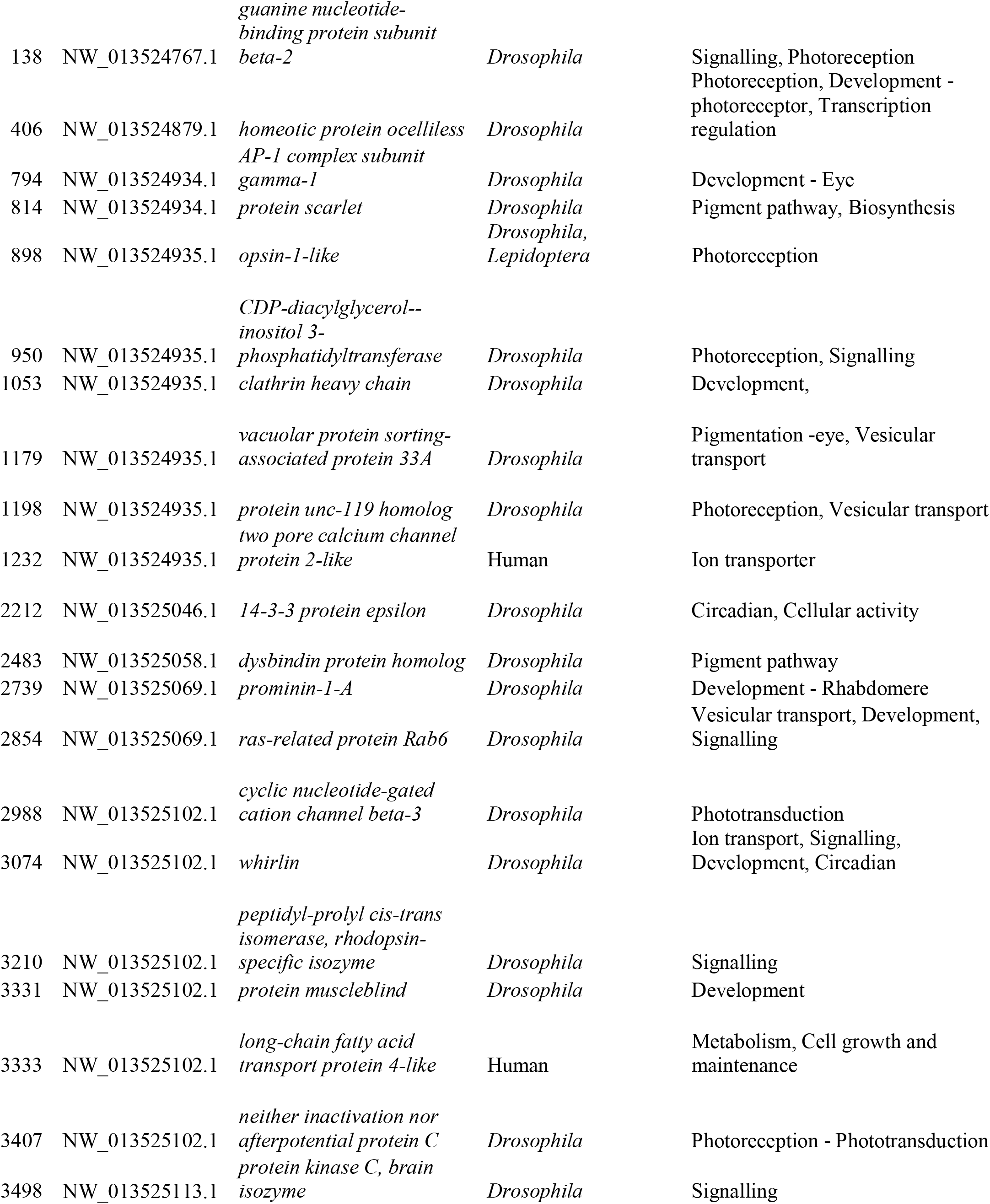
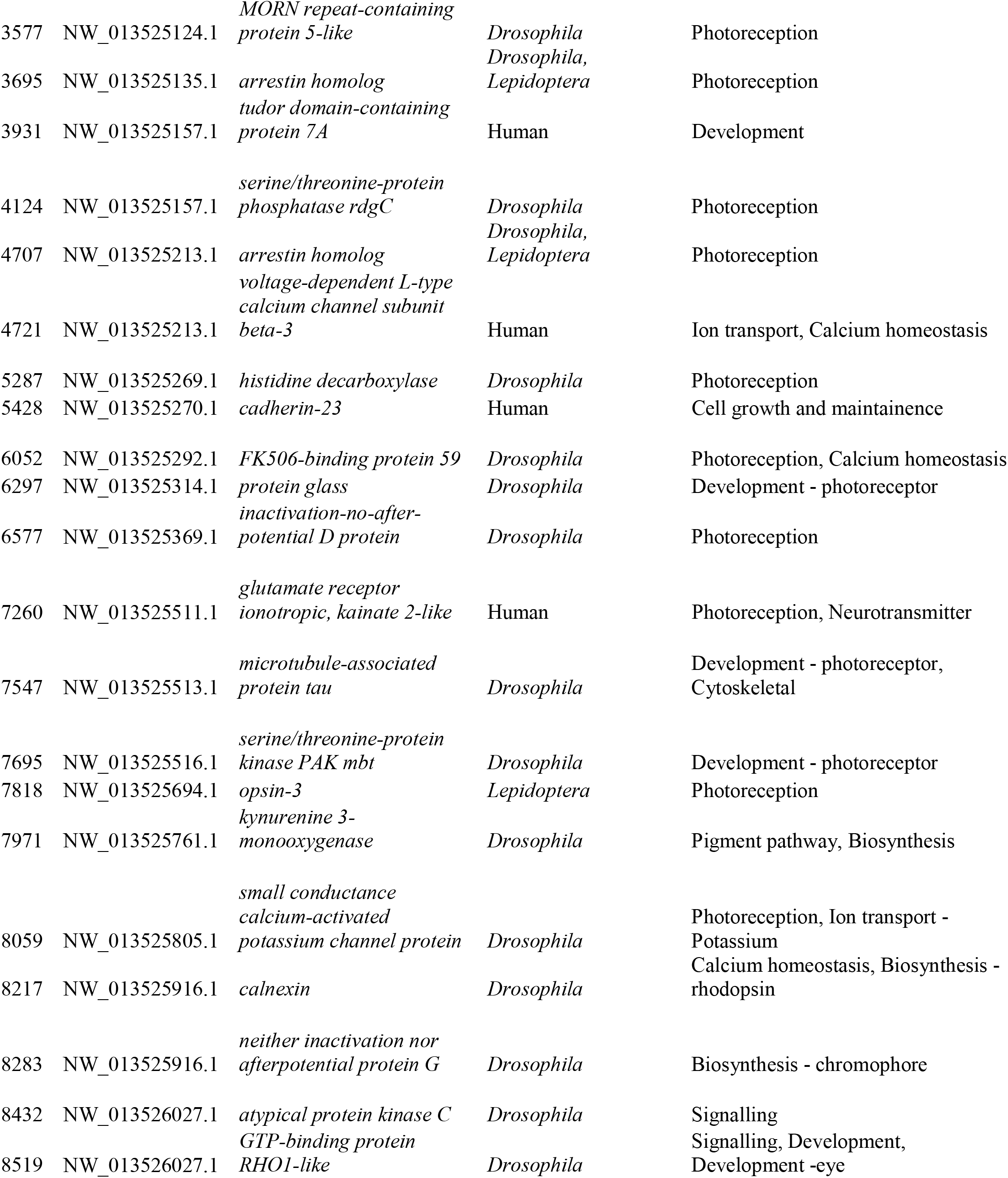
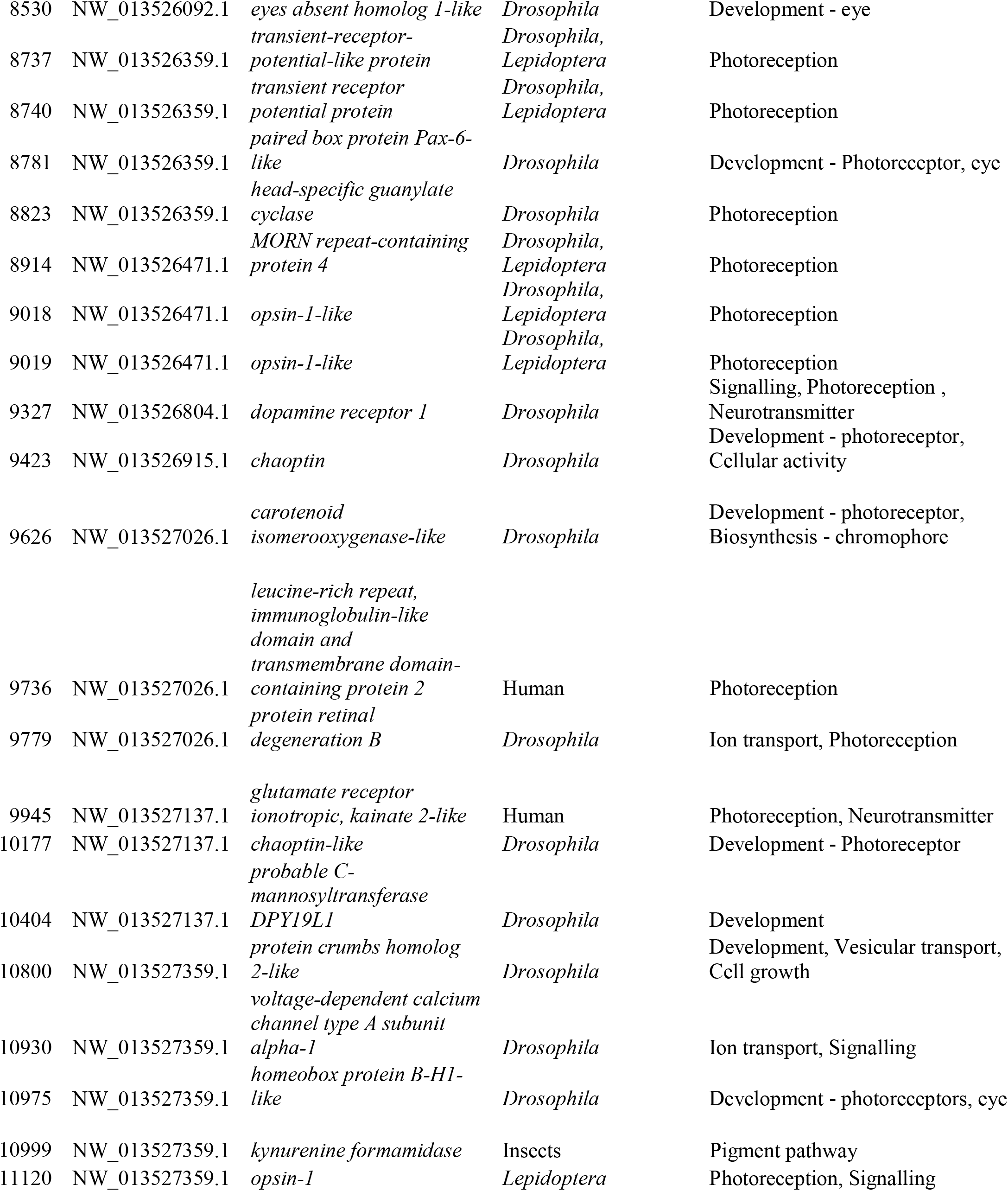
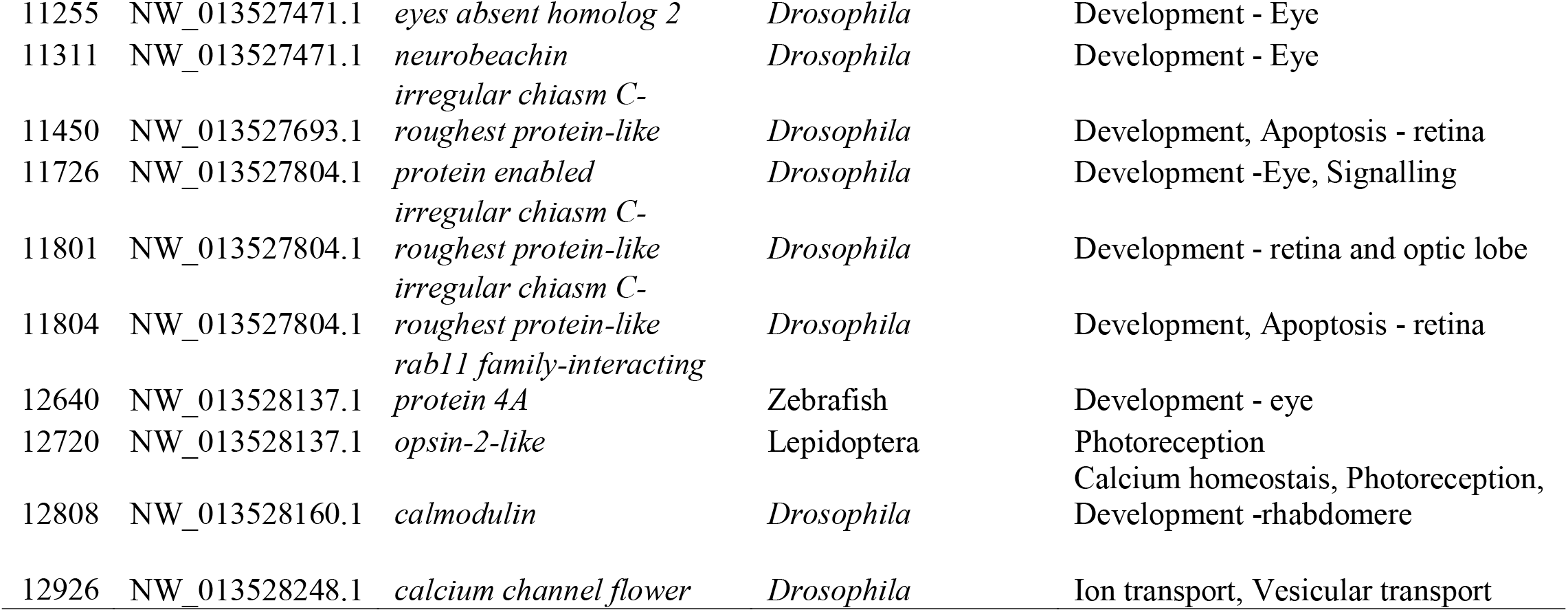
Functional annotation of genes showing upregulated expression in eyes.

**Table S8: Sample details of sequences and genomes used in the study.** See Excel file “TableS8_SpeciesListAndGenomeDetails.xlsx” for species details, accession numbers and genome IDs from Genbank, Lepbase and GigaDB.

**Table S9: Substitutions in amino acid residues within 5 Å distance of the chromophore binding site.** See excel file “TableS9_ResiduesWithin5Å.xlsx”

**Figure S1.**
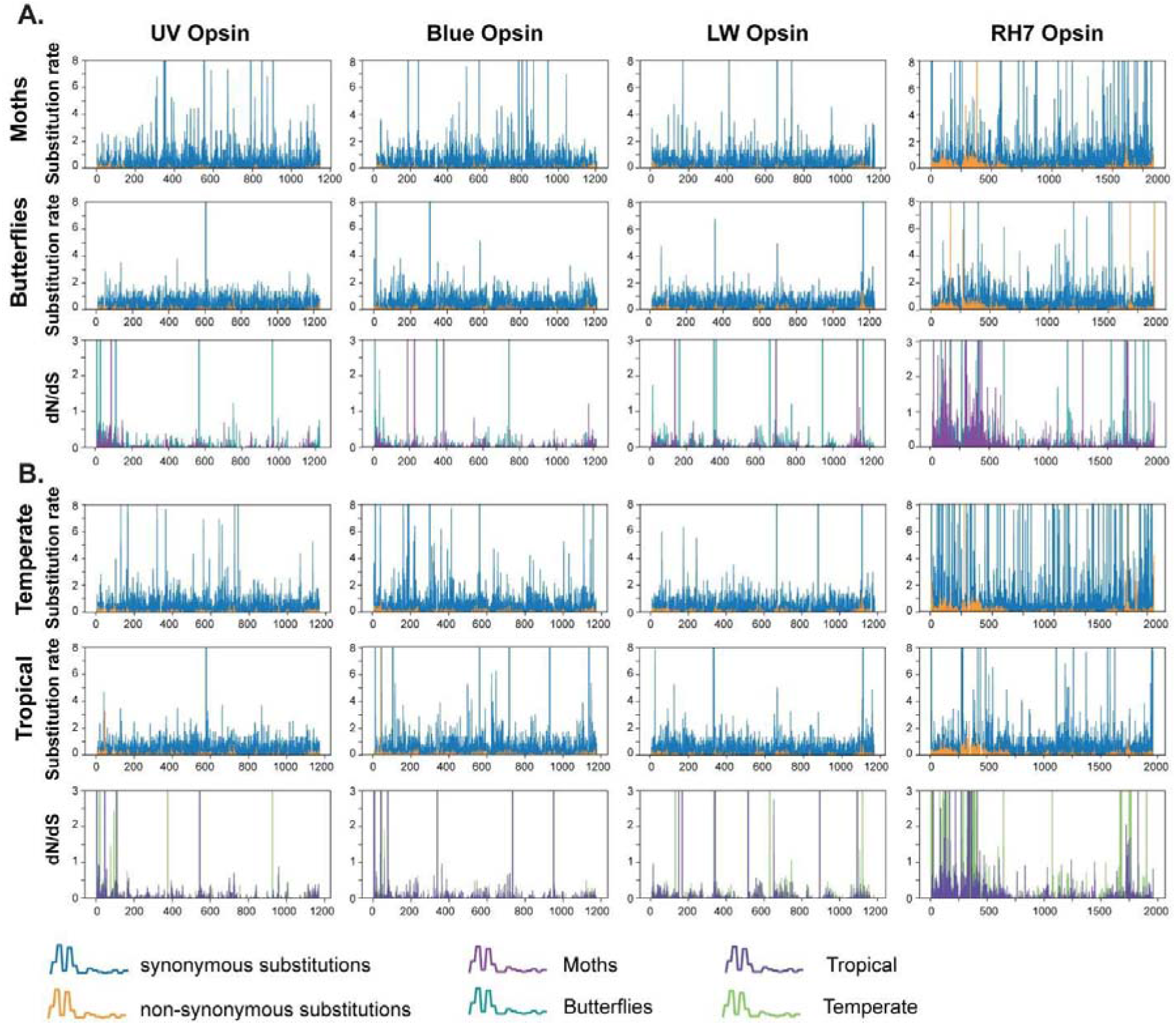
Comparative molecular evolution of opsins in Lepidoptera from i) different climate zones ii) Lepidoptera occupying separate diel-niche. A. Comparison of rates of molecular evolution between butterflies and moths is shown. First two panels represent nonsynonymous (orange) and synonymous (blue) substitution rates for each codon while the third panel represents dN/dS ratio for butterflies (pink) and moths (turquoise). B. Molecular evolution rates between temperate and tropical Lepidoptera are shown with synonymous (blue) and nonsynonymous substitution rates (orange) for each codon (first and second panel) as well as dN/dS ratios for tropical (purple) and temperate (green) are shown (third panel)

**Figure S2.**
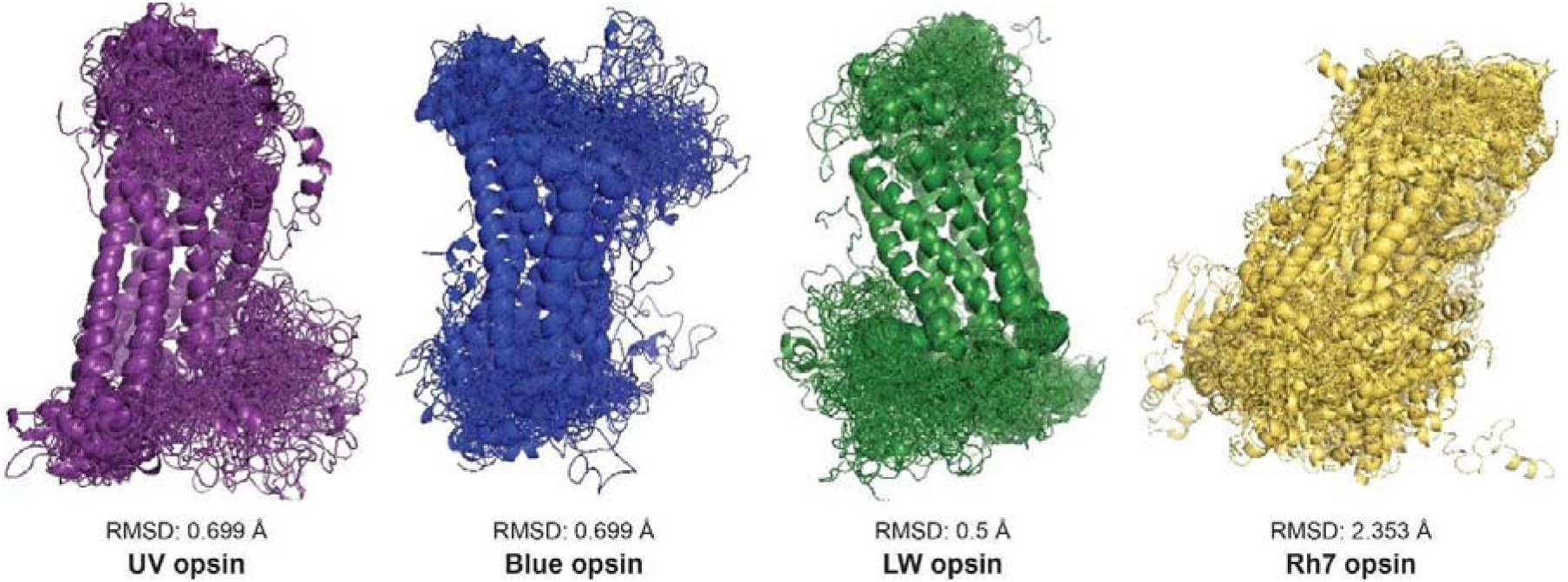
Structural resemblance of opsin proteins by superimposition of entire protein structures.

## Notes

### Competing Interest Statement

The authors have declared no competing interest.

